# Dissecting the regulatory and genomic drivers of the dichogamy determining G-locus in pecan

**DOI:** 10.64898/2026.05.14.725045

**Authors:** John T. Lovell, Hormat Rhein, Avinash Sreedasyam, Nolan Bentley, Patricia Klein, Warren Chatwin, Lillian Padgitt-Cobb, Avril M. Harder, Paul Grabowski, Sarah B. Carey, Alex Harkess, Jerry Jenkins, Chloee M. McLaughlin, Christopher Plott, Joanna Rifkin, Joe Song, Jenell Webber, Melissa Williams, Jane Grimwood, Jeremy Schmutz, L.J. Grauke, Xinwang Wang, Jennifer J. Randall

**Affiliations:** Genome Sequencing Center, HudsonAlpha Institute for Biotechnology, Huntsville, AL 35806, USA; U.S. Department of Energy Joint Genome Institute, Lawrence Berkeley National Laboratory, Berkeley, CA 94720, USA; New Mexico State University, Las Cruces, NM 88003, USA; University of Texas at Austin, Austin, TX 78712, USA; Texas A&M University, College Station, TX 77843, USA; USDA-ARS, Crop Germplasm Research Unit, College Station, TX 77845, USA

## Abstract

Many hermaphroditic species increase outcrossing rates by partitioning reproduction so that male and female organs mature at different times, a phenomenon known as dichogamy. Previous work has documented that dichogamy in pecan trees is governed by the Mendelian super-gene “G-locus”; however, its non-recombinant sex chromosome-like architecture has impeded quantitative genetic exploration and candidate gene discovery. Here, we probe the genetic drivers of the G-locus through a pangenome-integrated quantitative genomics experiment. We first provide one of the clearest examples to date of mapping bias, where a linear reference-based GWAS discovered 66 off-target peaks while mapping with a pangenome graph reference resolved the known single Mendelian locus. Across six new genome assemblies, the fully haplotype phased G-locus QTL spanned 223-491kb and included 25 candidate gene families. The strongest candidate gene encoded a MATE efflux protein and had dominant allele-specific action during male flower developmental stages. Combined, these candidates and genomic resources provide a powerful foundation for breeding and optimal dichogamy phenotype engineering for future pecan orchards.

## INTRODUCTION

Dichogamy, the temporal separation of stigma receptivity and mature pollen shedding, is a widespread mechanism to increase outcrossing in flowering plants (Lloyd and Webb 1986; Bertin and Newman 1993; Barrett 2003). While valuable in wild populations to maintain genetic diversity and increase heterozygosity, heterodichogamous populations, which segregate dichogamy phenotypes, can present major challenges for breeders and orchard managers (Grauke and Thompson 1993; Michler et al. 2004) requiring the use of pre-determined pairs of heterodichogamy compatible cultivars that allow for successful pollination and nut production.

Pecan trees (*Carya illinoinensis* (Wangenh.) K. Koch) are monoecious diploids that manifest dichogamy as protandrous (type I, male-first) and protogynous (Type II, female-first) flowering patterns. Previous work has demonstrated that heterodichogamy in pecan is controlled by a single Mendelian ‘G’-locus (Thompson and Romberg 1985), where *gg* homozygous recessive and *Gg* heterozygous individuals display protandrous and protogynous phenotypes respectively. Importantly, *GG* protogynous individuals are known (Thompson and Romberg 1985; Bentley et al. 2019) and do not appear to suffer obvious deleterious effects. As such, unlike many sex determination systems, the molecular mechanisms that induce male and female floral function in pecan must be fully intact in both *G* and *g* haplotypes, and the genetic mechanisms that cause dichogamy segregation in pecan are very likely to be regulatory in nature.

Divergence between the two G-locus haplotypes predates the diversification of pecan and perhaps even the *Carya* genus (Groh et al. 2025). Given this divergence, and possible other genomic complexities typical of sex-determining regions (Hobza et al. 2015; Papadopulos et al. 2015; Carey et al. 2021), a single-haplotype reference genome is unlikely to accurately capture the genetic diversity present among G-locus alleles. Here resolved these issues and probe the genetic basis of dichogamy in pecan by (1) building a pangenome with eight (seven new to this work) genome assemblies that span both *G* and *g* haplotypes, (2) remapping the G-locus against the graph reference using the largest GWAS population for dichogamy explored to date, (3) describing the structure and genome evolution of the G-locus, and (4) analyzing allele-specific gene expression across a floral tissue development time course to highlight several genes with potential regulatory roles in dichogamy.

## RESULTS AND DISCUSSION

### Genome-wide associations with heterodichogamy

To test the hypothesis that the G-locus is the primary driver of the Mendelian inheritance pattern of pecan dichogamy, we estimated binary dichogamy type breeding values (typeI: protandrous - male first; typeII: protogynous - female first) for 121 genotypes in a single common garden. Importantly, this panel includes samples from four of the five main genetic subpopulations of native pecan (Grabowski et al. 2025) and from both cultivars and unimproved populations (Fig. 1A; Supplementary Data 1). As expected from a heterodichogamous species, protandrous and protogynous phenotypes segregated near 1:1 (42.1% protandrous, binomial *P* = 0.101). This segregation ratio is also consistent among breeding populations (progeny from experimental crosses = 48.3% protandrous, selected seedlings = 39.5%) and wild germplasm (40.7% protandrous; χ^2^ = 0.602, df = 2, *P* = 0.7401), indicating that bottlenecks associated with cultivation do not necessarily extend to the loci that drive heterodichogamy.

**Figure 1.**
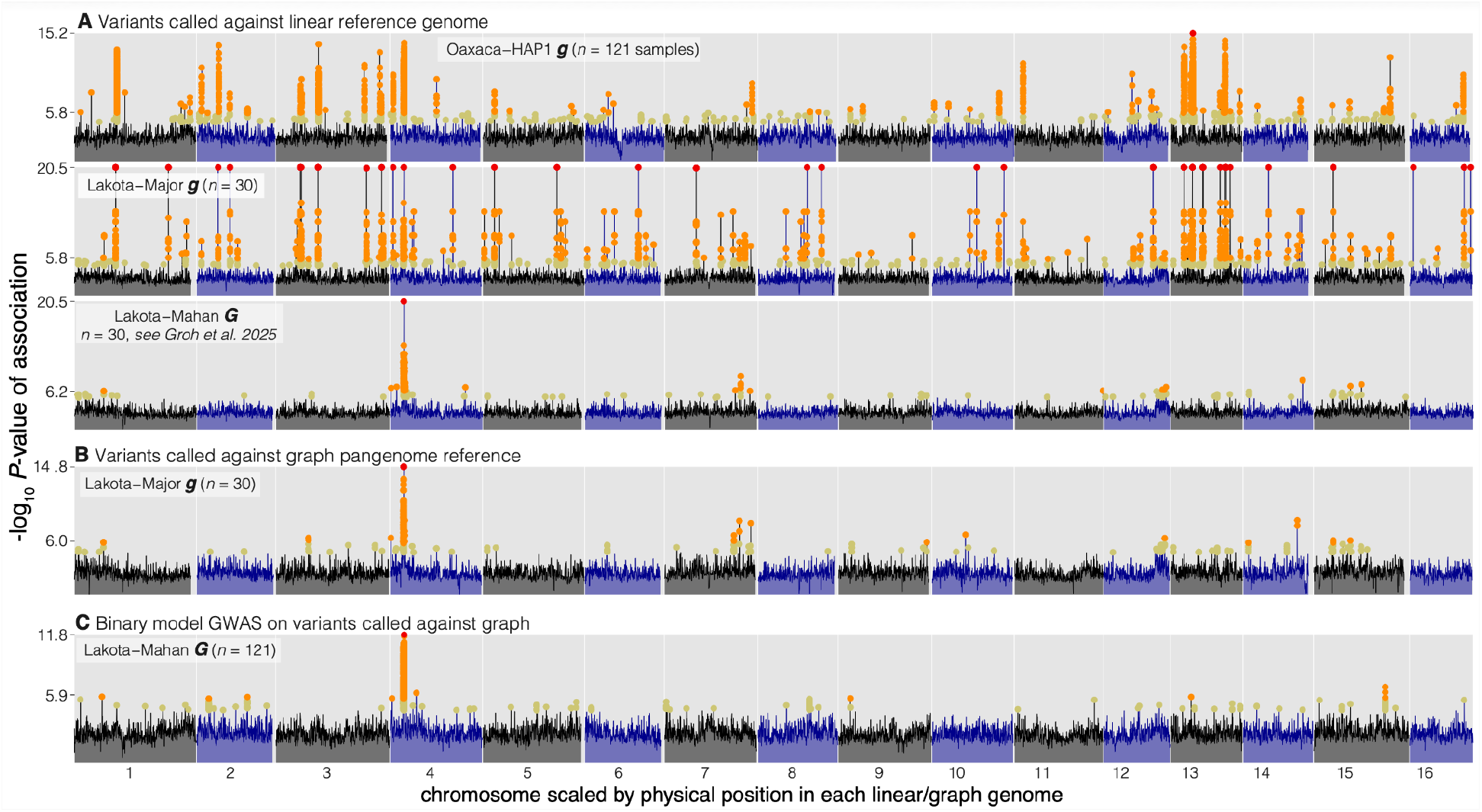
Genome-wide associations by reference genotype and structure. Variants were called against five references, grouped into linear (**A**) and graph pangenome (**B-C**) structures. In all panels, the maximum −log_10_ *P*-value association within non-overlapping 100kb intervals is plotted genome-wide (black/blue area plots) and all significant associations are plotted as colored points (red: most significant, orange: FDR *P*-value < 0.005, yellow: FDR *P*-value < 0.05). To be consistent with methods of Groh et al. (2025), the top four GWAS scans were run using a linear model in GEMMA treating the dichogamy trait as continuous. All *n* = 30 scans use exactly the same samples as Groh et al. (2025). **C** This scan was run with a more statistically appropriate binary model in LDAK using all 121 samples phenotyped here.

Our initial genome-wide association mapping (GWAS) scan using variants called against the existing Oaxaca-HAP1 V3 linear reference genome (Grabowski et al. 2025) revealed that the G-locus on Chromosome 4 was by no means the only significant association (Fig. 1A). Instead, we observed 1,408 ‘off-target’ highly significant (FDR-corrected *P* < 0.005) associations outside of the G-locus spanning 66 unique peaks and 135.7kb of genomic sequence (Supplementary Data 2-3). To confirm that the off-target peaks were not an artifact of GWAS methods, we extracted 21bp ‘diagnostic’ sequences that were fixed present-absent between short read libraries of six *GG* and *gg* individuals. These 21-mers masked 82.2kb of sequence across 2,632 unique positions outside of the g-locus (39.4% not masked by 21mers; Supplementary Data 4), providing further support that sequence outside of the G-locus segregates perfectly with dichogamy phenotypes and G-locus haplotypes.

The abundance of strong off-target signals comes in sharp contrast to the single locus peak observed by Groh et al. (Groh et al. 2025) (hereon ‘Groh’). We hypothesized that the difference in GWAS topology between these scans were methodological artifacts attributable to differences in reference G-locus haplotypes. Specifically, Groh mapped to the LakotaV1 genome, which contained the dominant *G* haplotype, while the Oaxaca-HAP1 used above had the recessive *g* haplotype. Despite its established role as a driver of dichogamy, both the complexity of the G-locus and divergence between the two major haplotypes (dominant ‘*G*’ and recessive ‘*g*’) challenges traditional quantitative trait mapping and candidate gene discovery. For example, missmapping, especially for sequences private to the haplotype not represented by the reference sequence, can cause conspicuous off-target sites to also appear in genome-wide scans (Bentley et al. 2019).

To test this hypothesis that the GWAS peaks outside of the G-locus are methodological artifacts, we aligned the 30 resequencing Groh libraries to our two new ‘V4’ haplotype-phased Lakota genome assemblies. Importantly, the Lakota cultivar is heterozygous (*Gg*) at the G-locus and the two haplotypes are represented in separate assemblies (*G*: ‘Lakota-Mahan’, *g*: ‘Lakota-Major’). Comparisons between variants called against each haplotype clearly confirmed an artifactual basis of off-target GWAS peaks (Fig. 1A): we observed the same strong single G-locus peak as Groh when mapping to the dominant *G* Lakota-Mahan haplotype; however, we find 29 off-target peaks spanning 403 markers and 66.1kb with perfect SNP-trait associations when mapped against the recessive *g* Lakota-Major haplotype (Fig. 1A, Supplementary Data 2). Simply mapping to a closely related genome, but one with a vastly different key haplotype, completely resolved the single dichogamy-associated QTL.

### Pangenomics resolves severe reference biases

While there was no obvious single cause of the off-target peaks, Lakota-Major (*g*) off-target GWAS intervals colocalized with a conspicuous enrichment of both annotated repeats and 21mers that were found in the Lakota-Mahan (*G*) G-locus haplotype. For example, when compared to 10,000 random genomic intervals with the same size distribution, the 70 off-target peaks from alignments to the linear Lakota-Major reference have > 3-fold more occurrences of the 21mers (Wilcoxon W = 357,635, *P* = 0.0013). This enrichment potentiates issues related to inaccurate variant detection (i.e., ‘pseudoheterozygosity) from linear reference genome alignments. Further supporting this observation, we were able to resolve mapping biases by choosing a reference genome that contained the larger dominant (*G*) haplotype. However, this choice required knowledge of the causal genomic interval and reference haplotypes. Thus, selecting a ‘correct’ reference genome may not be a tractable solution to solve similar problems broadly across various phenotypes and genetic architectures. Pangenomics offers a potential generalizable solution to resolve mapping bias: by including multiple haplotypes in a ‘graph’ reference, it is possible that biases could be resolved without *a priori* knowledge of causal loci.

To test whether pangenomics solves reference biases when mapping pecan heterodichogamy, we built graph pangenome references using the previously published V3 Oaxaca-HAP1 (G-locus haplotype = *g*) & -HAP2 (*g*) references (Grabowski et al. 2025), and six new assemblies: previously mentioned Lakota-Mahan (*G*) & -Major (*g*), Pawnee-Mohawk (*g*) & -SHG (*g*), and Elliott-HAP1 (*g*) & -HAP2 (*G*, see methods and Supplementary Note 1 for details on genome assembly). To test generalizability regarding the G-locus haplotype, we separately constructed graph pangenomes with either the Lakota-Major (*g*) or Lakota-Mahan (*G*) assembly as the ‘primary’ reference in the Minigraph-Cactus pipeline (Hickey et al. 2023). Using both graph pangenome references, where the ‘primary’ reference differed in G-locus haplotype, we generated sets of variants and ran genome scans for the dichogamy phenotype. While some minor differences existed between the resulting graph reference GWAS scans, regardless of the primary reference, a graph pangenome resolves reference bias and pinpoints the expected single G-locus peak (Fig. 1B-C) without a resulting loss of power or precision.

This comparison between graph and linear alignments also provided some evidence about the proximate causes of off-target peaks. Using two representations of variants (graph and linear) anchored to the Lakota-Major (*g*) coordinate system, we probed how genotype calls differed for off-target peaks from dichogamy genome scans. Genotype calls from protandrous (*gg*) libraries were 92% concordant homozygous reference in both the linear- and graph-based results. However, nearly half (46%) of linear-called heterozygous genotypes in protogynous libraries (*Gg, GG*) were incongruous and called homozygous reference using the graph (Supplementary Figure 1). This form of miscalled genotyping is referred to as ‘pseudoheterozygosity’ (Jaegle et al. 2023). While perhaps not extensible to all genetic mapping studies, these results add to a growing body of evidence that pangenome graph alignments can resolve cases of severe mapping biases, particularly those resulting from pseudoheterozygous calls (Miao et al. 2025; Kuster et al. 2026; Morris et al. 2026).

### Comparative analysis of G-locus causal haplotypes

Armed with the knowledge that pangenome-anchored GWAS is the most robust approach, we remapped all short read libraries from all 121 genotypes with dichogamy phenotypes to the Lakota-Mahan (*G*) anchored pangenome graph and ran GWAS using a statistically-appropriate binary trait model (Fig. 1C). We then intersected the GWAS hits with Lakota-Mahan (*G*) regions masked by G-specific 21-mers, which resolved the G-locus to 6,483,535-6,952,982bp on Chromosome 4, where a discrete block of strongly-linked loci were nearly uniformly associated with protogyny (Fig. 2A). To determine the reference position of the G-locus in all references, we aligned the 469,447bp dominant *G* Lakota-Mahan haplotype to the other seven genome assemblies. The resulting coordinates (Table 1) largely recapitulate previous work: the recessive *g* haplotype is roughly half the size of the dominant and there is only moderate haplotype size variation within each class.

**Table 1.** G-locus positions across eight pecan genomes. The GWAS- and kmer-defined G-locus interval in the Lakota-Mahan reference genome was extracted and aligned to chromosome 4 of the other seven genomes via minimap2 -xasm20. *coordinates for Lakota-Mahan were defined by GWAS and 21mers. **The two Oaxaca haplotypes are identical by descent.

| G-hap. | Chr. | start | end | width | strand | Target genome |
| --- | --- | --- | --- | --- | --- | --- |
| *g* | Chr04 | 6765360 | 7012431 | 247071 | + | Oaxaca_HAP1 |
|  | Chr04 | 6775747 | 7022818 | 247071** | + | Oaxaca_HAP2 |
|  | Chr04 | 6541262 | 6801341 | 260079 | + | Elliot_HAP1 |
|  | Chr04 | 6529974 | 6787778 | 257804 | + | Lakota_Major |
|  | Chr04 | 6493328 | 6748223 | 254895 | + | Pawnee_Mohawk |
|  | Chr04 | 6698590 | 6922240 | 223650 | + | Pawnee_SHG |
| *G* | Chr04 | 6483535* | 6952982* | 469440 | + | Lakota_Mahan |
|  | Chr04 | 6492239 | 6982974 | 490735 | + | Elliot_HAP2 |

**Figure 2.**
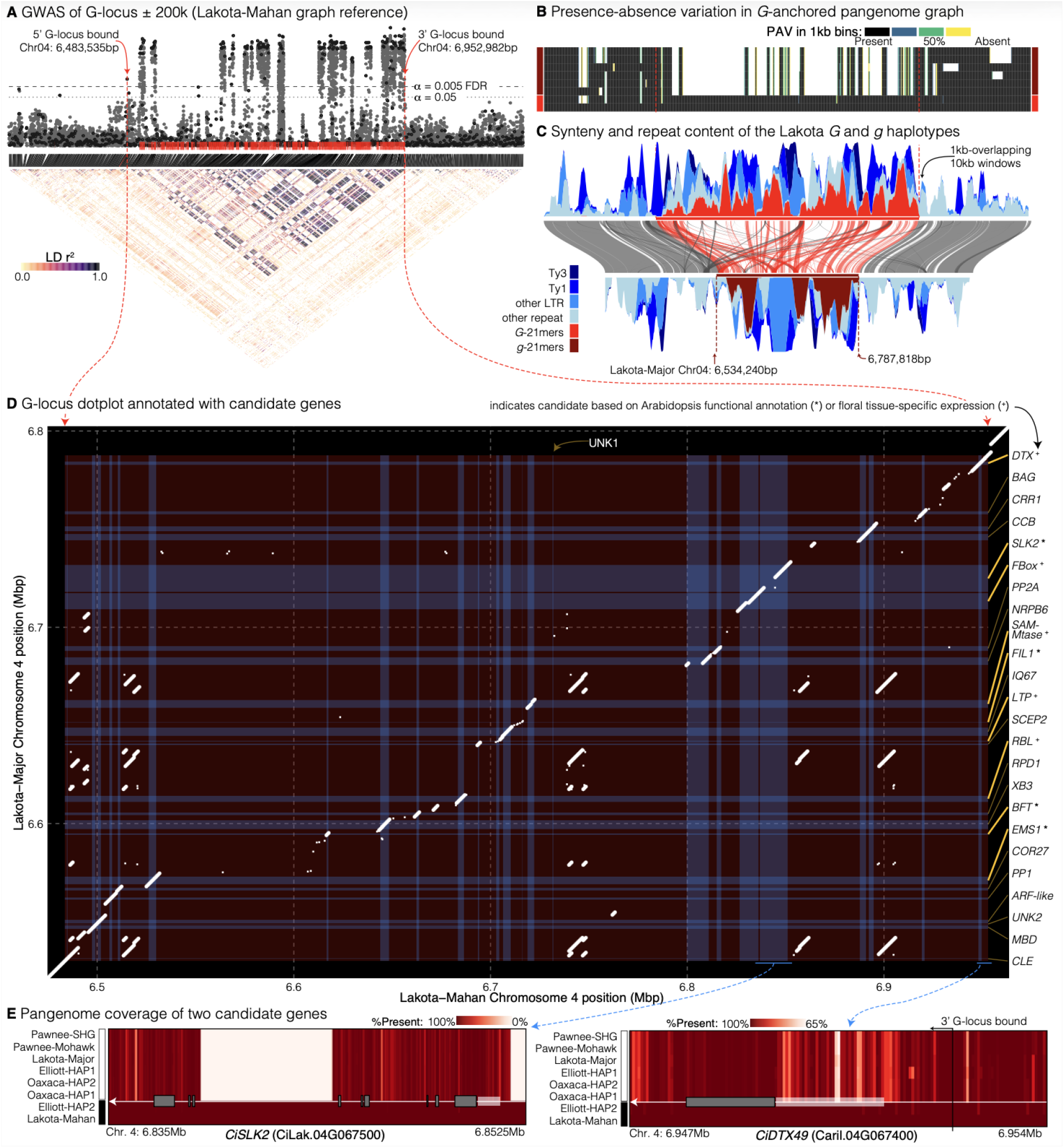
Inheritance and structural variation of the G-locus. **A** Binary Wald-statistic association −log_10_ *P*-values (range = 0.0-11.8) for the Lakota-Mahan G-locus ± 200kb and a linkage-disequilibrium heatmap. For visualization purposes, only one marker (dark grey points) every 1kb was plotted in the LD map. The chosen marker was either a random sample (if min *P*_FDR_ > 0.05) or the most significant hit. **B** Sequence presence-absence variation was calculated as the proportion of sequence traversed by each haplotype path in 1kb windows across the same interval as panel A; haplotypes from top to bottom: Oaxaca-HAP1, -HAP2, Pawnee-Mohawk, -SHG, Elliott-HAP1, Lakota-Major, Elliott-HAP2 and Lakota-Mahan. **C** For each window in both Lakota-Mahan (top area plot) and Lakota-Major (bottom area plot) sequence content was hierarchically assigned to four repeat categories then *G-* and *g-*21mers. In both plots, the y-axis range is defined by the dashed interval bound lines and scaled from 0-100%. Syntenic blocks (see panel D) connect the two haplotypes. Any blocks with both query and target sequences in the G-locus are red. **D** Nucmer alignment coordinates of the two G-locus haplotypes ± 20kb (white segments: ≥1kb alignment length, white dots = length < 1kb). The transcript coordinates of all G-locus genes in both references are annotated as vertical (Lakota-Mahan) or horizontal (Lakota-Major) blue rectangles, ending at the G-locus bounds for each genome. There is one PAV gene, “UNK1”, which is only found in Lakota-Mahan and labeled along the horizontal axis. High-priority candidate genes, based on their functional annotation (*) or tissue-specific expression in Arabidopsis (^+^) are flagged and have thicker yellow line segments connecting the label. **E** Sequence presence-absence variation at *CiSLK2* (left), which has two large sequence deletions in an intron and just upstream, and *CiDTX49* (right), which has highly conserved coding sequence but significant divergence in the upstream regulatory region.

Like a sex chromosome or self-incompatibility locus (Kamau and Charlesworth 2005), the expectation is that the G-locus super-gene contains multiple causal loci that need to be in linkage phase to ensure proper functioning of the molecular machinery that induces dichogamy (Thompson and Jiggins 2014). The resulting linkage dissequilibrium (LD) block was evident across the entire population (Fig. 2A), although LD was not as strong among homozygous *gg* recessive individuals (Groh et al. 2025). Perhaps as a consequence of recombination suppression, fixed sequence insertions in the dominant *G* haplotype, or equivalently, sequence deletion in the recessive *g* haplotype, have caused >200kb differences in haplotype sizes (Fig. 2B). Many of the large blocks of sequence only found in the G-haplotype are composed of sequence enriched in EDTA-annotated repeats (Fig. 2C) or multi-copy syntenic blocks (Fig. 2D, Supplementary Data 5). This expansion of repetitive elements is often attributed to a relaxed evolutionary constraint via suppressed recombination and is common in the dominant haplotype of plant sex determination loci (Kejnovsky et al. 2009; Hobza et al. 2015), similar to the mammalian Y or avian W chromosomes (Ezaz and Deakin 2014).

Since selection should favor elements that reduce recombination in such sex-linked loci (Charlesworth and Charlesworth 1973; Jay et al. 2024), it is somewhat surprising that large inversions were never observed between *G* and *g* haplotypes or the surrounding interval (Fig. 2C-D). However, there were many minor repeat-associated translocations particularly at the 5’ bound of the G-locus (Fig. 2D). When combined with the massive array of large sequence insertions/deletions, these translocations could both be a cause and result of low recombination throughout the G-locus.

### Positional candidate genes for dichogamy

Given its known inheritance and segregation, causal dichogamy genes must be physically present in the G-locus and contain (1) fixed presence-absence (PAV), (2) protein-coding, or (3) regulatory variants that segregate between allele classes. To reduce the potential for PAV artifacts (Brůna et al. 2026), we used a direct lift-over approach (see methods for details) to structural gene annotation and projected representative (both *G* and *g* alleles) gene models from the V1 genome annotations onto all eight genome assemblies (Supplementary Data 6).

We defined 24 sets of orthologs that were constitutively present in the G-locus of all genomes (Fig. 2D, Table 2). This list included some of the same functionally interesting G-locus genes (annotated in figures and tables with a ‘*’) as other studies described: pecan orthologs to *SEUSS-LIKE KINASE2* (“*CiSLK2*”, (Lee et al. 2014)), *BROTHER OF FLOWERING LOCUS T* (“*CiBFT*”, (Pin and Nilsson 2012)), the anther-specific *EXCESS MICROSPOROCYTES1* (“*CiEMS1*”, (Zhao et al. 2002)), and a gene similar to the stamen-specific floral regulator *FIL1* (“*CiFIL1-like*”, (Nacken et al. 1991; Groh et al. 2025)). Arabidopsis homologs of four other candidate genes in the interval have conspicuous floral tissue-specific expression (Fucile et al. 2011) in Arabidopsis (annotated in figures and tables with a ‘^+^’), including stamen-specific MATE efflux family protein gene, *DETOXIFICATION 49* (“*CiDTX49*”), that had not previously been highlighted as a candidate causal gene.

**Table 2.**
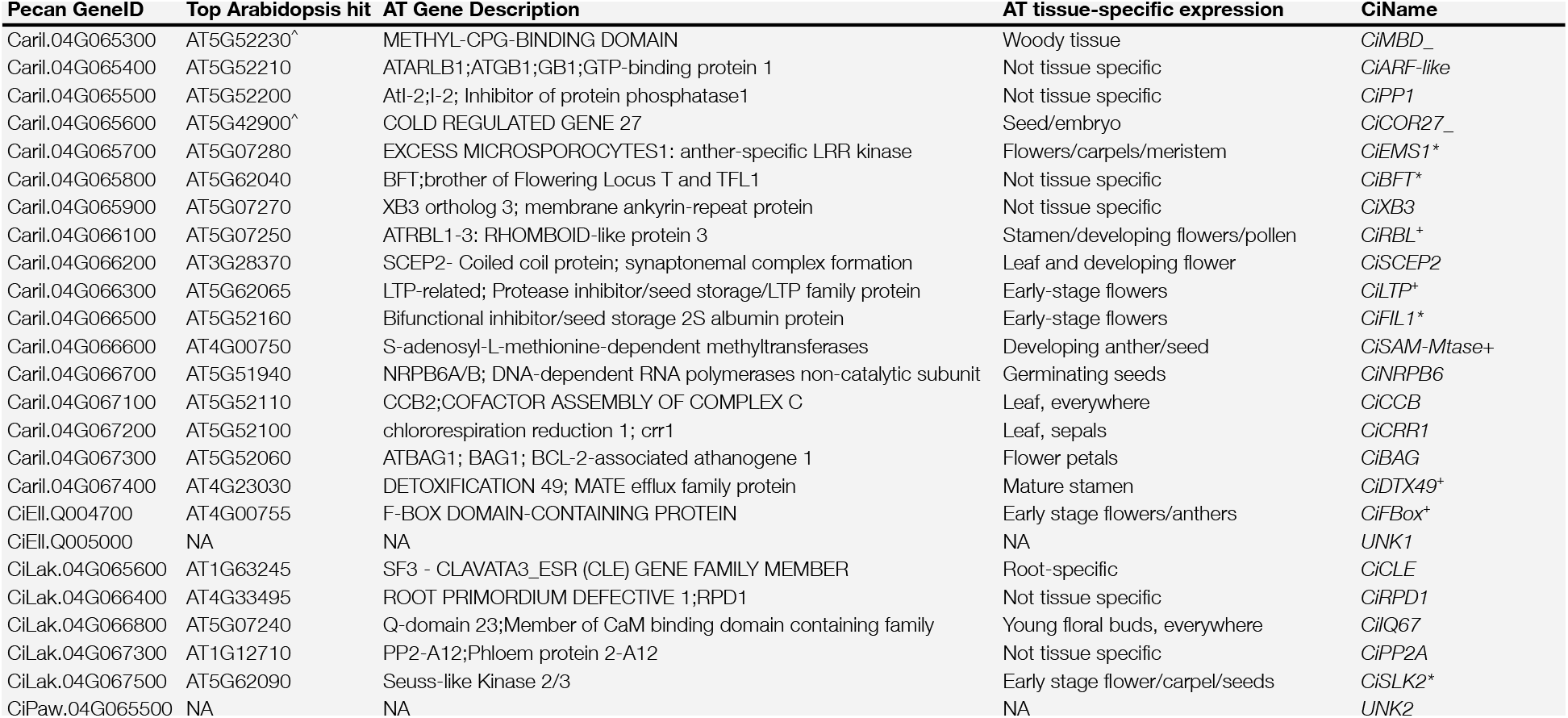
Candidate genes in the G-locus. List of 25 candidate gene families in the G-locus. For each Carya V1 annotation name (first column), the top Arabidopsis thaliana blast hit and functional information are reported. Summary of tissue-specific expression (Fucile et al. 2011) are also included. To improve interpretation, we typically refer to each gene its “CiName”, which uses the top Arabidopsis homolog ID (or functional description). Where appropriate, “*”, “^+^” and “_” are appended to the CiName to indicate strong functional candidates, genes with floral tissue-specific expression, or low blast support (flagged with “^^^” in the top Arabidopsis hit column), respectively. Genes with no significant BLAST hits are named “UNK”.

Unlike most sex determination loci in other taxa and in contrast to its genomic sequence alignments (Fig. 2B-C), the pecan G-locus had remarkably low levels of gene PAV, and none of the top candidates exhibited copy number variation. Only one gene was annotated in a large sequence insertion/deletion (Fig. 2D): *G*-specific CiEll.Q005000 (hereon ‘UNK1’), a short single-exon gene without sequence similarity to genes in any plant species annotated on NCBI.

In contrast to the limited PAV, we found substantial putatively functional protein-coding variation among alleles of several candidate genes. The most striking example comes from *CiSLK2*, which, despite having largely syntenic alignments (Fig. 2D), had 74 coding SNPs and 24 INDELs between the two ‘Lakota’ haplotypes and large sequence deletions both immediately upstream and in an intron (Fig. 2E). The limited gene PAV in the G-locus is perhaps expected because, unlike most sex chromosomes, rare GG individuals (e.g. ‘Mahan’ and ‘Apache’, (Bentley et al. 2019)) are known to exist in breeding populations and can produce viable gametes, so complete loss of necessary sex-specific genes, as is common on the dominant sex determining allele in other systems, is not likely to have occurred in the pecan G-locus. Instead, regulatory variation, like that observed extensively in the upstream region of *CiDTX49* (Fig. 2E) is the most probable causal mechanism.

### Tissue-specific expression across G-locus alleles

To explore the impacts or regulatory variation, we conducted an RNA-seq time course experiment (Supplementary Data 7-8) across six pecan cultivars, two of each G-locus genotype class: ‘Pawnee’ & ‘Major’ (*gg*), ‘Lakota’ & ‘Elliott’ (*Gg*), and ‘Mahan’ & ‘Apache’ (*GG*). Two biological replicates of all genotypes were grown in the USDA pecan repository orchard and total RNA was extracted and sequenced from four tissues: (1) dormant bud, (2) swollen bud, (3) immature pistillate flower, (4) immature catkin. These tissues encompass the full range of regulatory element effects as the buds transition from dormancy to mature male and female flowers.

Overall, tissue-specific total transcript abundance dovetailed with observations in other systems. The two clusters of coexpressed female- and male-specific genes (top 6 rows Fig. 3A) were all homologs of Arabidopsis genes expressed or functionally annotated in developing flowers, with the exception of *CiCLE3* (homolog of Arabidopsis *Clavata3*), which was only expressed in one condition. For example, *CiFIL1-like* and *CiEMS1* were highly upregulated in catkins across four of the six genotypes, which aligns well with their known male-specific expression in Arabidopsis, and pecan homologs of *BFT*/*TFL* has been detected at high levels in pecan catkins (Rhein et al. 2023). However, it also appears that regulatory roles in pecan have diverged at some loci, like *CiDTX49*, which is highly upregulated in Arabidopsis anthers, but expression of *CiDTX49* was significantly upregulated in only in female developing pistillate tissue (Fig. 3A).

**Figure 3.**
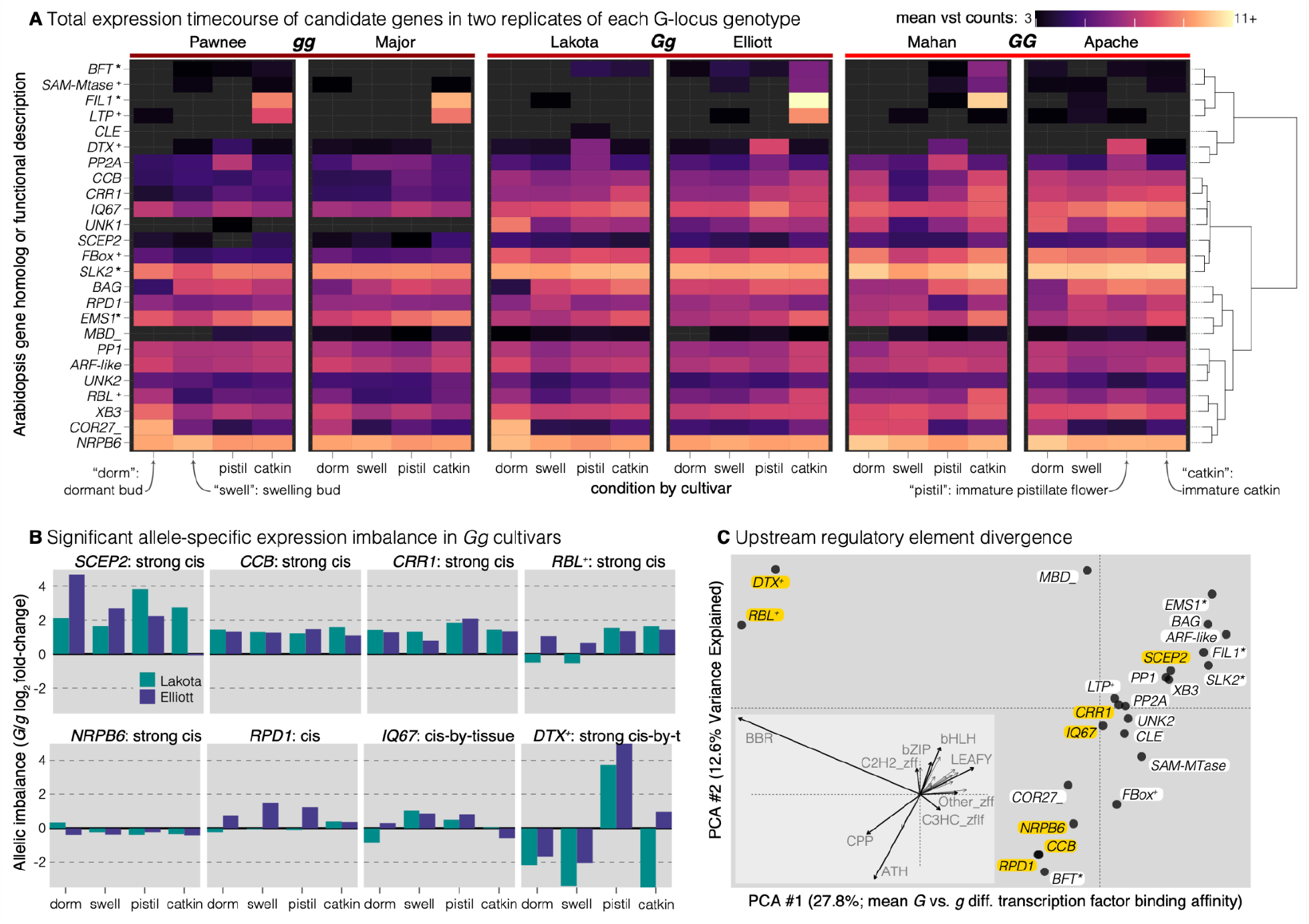
Total and allele-specific gene expression variation in the G-locus. **A** Mean Variance-Stabilized-Transformed total transcript abundance counts (empty if no reads mapped) clustered by between-gene coexpression. For visualization purposes, we cropped the heatmap scale to 11, but one sample exceeded this: *CiFIL1-like* catkin (VST = 16.3). **B** Allelic imbalance log_2_ fold-change (*G* vs *g*, positive: *G* > *g*) of eight genes with significant additive or tissue-specific cis regulatory element effects for Lakota (left/teal) and Elliott (right/purple). Effects are labeled as either constitutive (‘cis’) or depending on tissue (‘cis-by-tissue’) and flagged with ‘strong’ for genes with FDR-adjusted *P*-value < 0.001. **C** Differential transcription factor binding affinity of all 24 single-copy candidate genes, highlighting the six cis- and two cis-by-tissue regulated genes (gold label backgrounds). Principal components were calculated from gene-specific mean differential binding affinities of the 1kb upstream region for 805 putative transcription factor binding motifs, averaged for the 23 TF classes. In the inset, vectors from the origin indicate the PC axis rotations due to mean differential binding affinity, with nine of the least correlated classes labeled.

The expression time course also revealed potential roles of genes that have not previously been functionally annotated in floral development, including *CiLTP* (Caril.04G066300, Arabidopsis ortholog: AT5G62065) and *CiPPT2* (CiLak.04G067300, AT1G12710), which were conspicuously coexpressed with *CiFIL1-like* and *CiDTX49* respectively. Other *a priori* functional candidates like *CiSLK2* were consistently regulated across tissues and genotypes; functional dichogamy roles for genes like this most likely relate to protein sequence or allele-specific effects.

### Cis-regulatory evolution of candidate genes

Since we have mapped the causal locus to the physical bounds of the G-locus, functional regulatory alleles must be proximate to and linked with the target genes: cis-regulatory elements, ‘CREs’. The genomic resources built here provide an ideal foundation to test for the presence of CREs through the ‘composite’ test of allele-specific expression (Wittkopp et al. 2004). In short, if a target gene is regulated by only unlinked trans-acting factors, both alleles in heterozygous individuals will be acted on equally by the regulatory environment and thus expressed at identical levels. However, additive or dominant effects of CREs can act solely on the local haplotype (e.g. upstream promoter or enhancer elements) and are evident by biased expression of one allele over another within a heterozygote. To test for the role of CREs in dichogamy, we assayed allele-specific expression (ASE, Supplementary Data 9) via a competitive mapping approach (Lovell et al. 2018) and modeled cis- and cis-by-tissue effects across the RNA-seq time course via likelihood ratio test comparisons of hierarchical linear models.

We expected causal genes to have similar and strong ASE in both heterozygous cultivars (‘Lakota’ and ‘Elliott’) analyzed here. It was interesting to note that, despite strong differential total transcript abundances between tissue types, *CiFIL1-like* showed no evidence of allelic imbalance in either heterozygote or any developmental stage, and allelic imbalance of *CiBFT* was strong but variable depending on genotype and tissue (Supplementary Fig. 2). Expression levels of these and other genes without evidence of CREs could still cause differences in dichogamy phenotypes through functional sequence differences or downstream regulatory action; however, since both alleles are expressed equally between sequences, expression levels themselves are unlikely to be a causal mechanism.

Conversely, we observed strong constitutive cis-regulatory effects in several genes consistent with dominant action of the *G* allele (Fig. 3B). The strongest of these across-tissue cis effects were observed in *CiSCEP2, CiCCB*, and *CiCRR1*, each of which had >2 log_2_ fold-change of the dominant allele over the recessive. Of these, only *CiSCEP2* had observed floral tissue expression in Arabidopsis (Table 2). Further complicating matters, the upstream 500bp putative regulatory region of these genes held few clues for the mechanisms driving dominant allele-specific action: differences between the upstream *G* and *g* haplotypes of the three were characterized by generally unrelated signatures of differential binding affinities across 23 transcription factor classes (Fig. 3C). Despite a lack of functional connections with CREs, the dominant action of the dominant allele highlights potential functional roles for these genes.

Since heterozygotes harbor both alleles of the causal genes and express both male and female function, we expected the proximate causal genes to not only have cis-regulated ASE, but to also have stage-specific allelic imbalance. For example, the *G* allele may be expressed primarily during female-first floral initiation of the pistillate flower, while the *g* allele of a causal gene may be upregulated during the maturation of male catkins. Across all candidates, only two genes had statistically significant cis-by-tissue interactions (*CiDTX49* and *CiIQ67*) and only *CiDTX49* had the expected pattern of dominance-switching depending on tissue stage (Fig. 3B). Expression in *CiDTX49* was generally biased towards the recessive *g* allele, except in immature pistillate flowers where both Lakota and Elliott expression was very strongly biased towards the dominant *G* allele. Unlike the additive CRE candidates, differential transcription binding affinity of *CiDTX49* was clear and highly similar to *CiRBL*, a gene with borderline cis-by-treatment (FDR-adjusted *P* = 0.086) ASE imbalance (Fig. 3C). The 500bp upstream putative promoter sequence for both Lakota-Mahan and Elliott-HAP2 dominant *G* alleles of both genes had highly enriched affinity for the three motifs associated with *Barley B Recombinant/Basic PentaCysteine* (*BBR*/*BPC*) transcription factors. *BBR/BPC* TFs are known to drive floral transitions and underlie sex determination pathways in other systems (reviewed by (Charlesworth 2021; Sahu et al. 2023)). While not initially intuitive, it is not surprising that none of the 10 BBR/BPC genes or in fact any other transcription factors in the Lakota V1 genome are found within the physical bounds of the G-locus. Such trans-acting regulatory factors must be able to access both alleles so that differential binding affinities cause cis-regulated differential expression of causal genes.

## Conclusions

To date, despite positional verification of the G-locus, the molecular mechanisms controlling heterodichogamy in pecan remain largely unknown. Several studies have proposed candidate genes; for example (Groh et al. 2025) and (Rhein et al. 2023) suggested that nonsynonymous fixed differences between the *G* and *g* alleles and differential expression of transcription factors could drive dichogamy. Indeed, both types of variation are certainly likely to play a role in the differential development of male and female floral organs. We also observed a single presence-absence variant gene that is correlated with, but not necessarily causal for protogeny. The limited gene PAV observed in the G-locus here is perhaps expected because, unlike most sex chromosomes, rare *GG* individuals are known to exist in pecan breeding populations (e.g. ‘Mahan’ and ‘Apache’, (Bentley et al. 2019)). *GG* individuals are neither weak nor reproductively limited as *GG* ‘Mahan’ is one of the three major founder lines that has been used to develop commercial pecan cultivars by the USDA breeding program and is known for its large nut size and prolific yield. Therefore, a complete loss of necessary sex-specific genes, as is common on the dominant sex determining allele in other systems, is not likely to have evolved in the pecan G-locus.

The single-locus dominant action of the G-locus and temporally variable expression of both sexes in pecan trees intimates that neither large-effect protein coding variants nor trans-regulatory elements can alone cause dichogamy. Instead, the biology and genetic architecture of dichogamy strongly suggests a central role for dominant cis-regulatory evolution mechanisms that drive floral timing and thus dichogamy in pecan. Here, we observe evolved differences between the G-locus alleles in the promoter regulatory regions and within introns, including large insertion-deletion polymorphisms. For example, the upstream region of the strongly cis-regulated and tissue-specific gene *CiDTX49* has undergone large-scale replacement of transcription factor binding sites between the two G-locus alleles (Fig. 3D). However it is important to note that a lack of allelic imbalance does not rule out candidate genes. Development-associated gene expression is notoriously transient (Wellmer et al. 2006) and further inquiry into dichogamy-associated transcriptional networks is certainly needed. This is especially true among genes like the strong functional candidate *CiFIL1-like*, which retains several fixed large sequence deletions in potentially regulatory regions of the recessive *g* haplotype. Combined, the regulatory regions and large scale functional variants of several candidate genes analyzed here provide strong targets for further functional validation, future genome engineering, and ongoing genome-enabled breeding efforts.

## METHODS

### Genome Assembly

Genomes were assembled following our previously developed pipelines that have been optimized for pecan, with minor modifications (see Supplementary Note 1 for complete assembly methods for each reference haplotype). In short, PacBio libraries were sequenced on the Sequel II or Revio platforms. The resulting HiFi reads (38.6-113.6× coverage) were co-assembled with Dovetail Omni-C reads (63.9-183.6× coverage) using HiFiASM+HiC (Cheng et al. 2021), scaffolded with the Juicer pipeline (Durand et al. 2016) and polished with 52.5-56.9× coverage Illumina PCR-free 2×150bp paired end reads to correct homozygous SNPs and short indels. Where parental strains were known (Pawnee and Lakota cultivars), post-hoc phasing was conducted by trio binning with 36.7-64.1× coverage Illumina PCR-free 2×150bp paired end reads from the parental genotypes. It is important to note that the ‘Oaxaca’ genotype is known to be partially inbred (Grauke et al. 2011, 2016) and the two G-locus haplotypes are 100% identical by descent (Supplementary Fig. 3). Therefore, we avoided pseudoreplication and only used HAP1 for candidate gene screening.

### Annotation of protein-coding genes and repetitive elements

Repeats in all genome assemblies were annotated with EDTA (Ou et al. 2019). Representative coding sequences (CDS) were obtained from orthogroups identified using GENESPACE (Lovell et al. 2022) across four previously published *Carya illinoinensis* cultivars (87MX3, Elliott, Lakota, and Pawnee) (Lovell et al. 2021). A single representative CDS from each orthogroup was selected to ensure non-redundant coverage of conserved gene families and to avoid inclusion of multiple gene copies. These CDS sequences were aligned to the eight haplotype-resolved assemblies sing GMAP v2025-04-19 (Wu and Watanabe 2005), a splice-aware genomic aligner suitable for predicting exon–intron structures. Indexed GMAP databases were first generated for each phased genome, and alignments were performed using the parameters --no-chimeras -B 5 --format=2 --min-intronlength=9 --max-intronlength-middle=500000 --max-intronlength-ends=10000--fulllength --npaths=1, which constrained each CDS to its best genomic alignment without chimeric mapping. Resulting gene models were stringently filtered, retaining only alignments with ≥ 70% sequence identity, ≥ 70% query coverage, and a minimum predicted protein length of 29 amino acids. Gene models failing these criteria were excluded to minimize annotation false discovery. Importantly, since the V1 genomes spanned two *g* and two *G* haplotypes (both Elliott-V1 and Lakota-V1 primary assemblies were *G*), we were confident that most functional alleles were accurately captured in V1 annotations. Combined, this produced V2 annotations that were of identical completeness as the original V1 resources (BUSCO = 97.7-98.3; BUSCO_Table).

### Dichogamy phenotyping

The catkin and pistil flowers of mature trees growing in the USDA repository orchards located in Somerville and Brownwood, Texas, were carefully monitored throughout the spring of 2023. For each tree, the presence of catkins, the timing of catkin shed, and the period during which pistils were fully receptive were recorded in detail. These flowering stages were used to track the timing and pattern of the male and female flowering development and determine the dichogamy for each genotype. A tree was considered protandrous when catkins released pollen before pistils became receptive, and protogynous when pistils reached receptivity prior to the maturation and shedding of catkins (Thompson and Romberg 1985).

### Genome-wide association mapping

Genome-wide association studies (GWAS) were implemented with two different models, where the dichogamy phenotype was treated as either a continuous or a binary trait. For genome scans where dichogamy was treated as a continuous trait, we used a mixed linear model and included a relatedness matrix generated with default settings implemented in GEMMA (v.0.98.30) (Zhou and Stephens 2012). We additionally ran genome scans using LDAK-KVIK (v6.1-1) where the dichogamy phenotype was treated as a binary trait by including the argument “--binary YES” in commands (Hof and Speed 2025). LDAK-KVIK uses a three step process to test for a marker’s association to phenotype. The first step is to construct leave-one-chromosome-out (LOCO) polygenic scores (PGS) for estimation of a test statistic scaling factor (λ) and is recommended to be run on a subset of sites included in the association study. For this step, we used PLINK (v.1.90b6.12) and thinned our variant set to only include sites with a minor allele frequency (MAF) > 0.01, and to remove sites within 100 kb with a squared correlation above 0.5, resulting in 572,048 variants. For the next two steps, we used default settings. All genome scans with n = 30 genotypes used the exact same samples reported in Groh et al. (2025). For all association studies, variants included in the scan were filtered to be biallelic, have less than 50% missingness, and a MAF ≥ 0.05 (*n* = 121 samples in genome scans) or minor allele count (MAC) ≥ 2 (*n* = 30 samples in genome scans). Using these parameters, reported is the total number of filtered variants included in genome scans using a linear reference: Oaxaca-HAP1 (*n* = 121) 14,608,838; Lakota-Major (*n* = 30) 16,178,227; Lakota-Mahan (*n* = 30) 15,806,817 and graph pangenome reference: Lakota-Major (*n* = 30) 16,178,227; Lakota-Mahan (*n* = 121) 16,202,781.

### De novo identification of fixed sequence-dichogamy associations

To identify the dichogamy locus in the Latoka reference, we used whole-genome resequence data from 12 genotypes (6 *GG* libraries: IXIR, IXYS, IXWF, IXXA, IXSW, IXKE and 6 *gg* libraries: IXDQ, IYBB, IXWM, IXZG, IYAP, IYBA). We first filtered the data using Trimmomatic v0.39, with leading and trailing values of 3, sliding window of 30, jump of 10, and a minimum remaining read length of 40 (Bolger et al. 2014). We identified all *k*-mers per library using meryl v1.3 count (Rhie et al. 2020). We next identified *k*-mers specific to the *gg* or *GG* genotypes. To accomplish this, we identified all 21-mers shared in *gg* genotypes using meryl’s intersect function, followed by a difference to identify the 21-mers never found in *GG* genotypes (and vice versa to generate a *GG*-specific k-mer list).

### Defining dichogamy associated peaks

The physical positions of all GWAS associations were processed independently for each reference genome, method, and sample set run combination. For each run, an ‘initial’ peak was defined as any set of highly significant hits (FDR *P* <= 0.005) with no gap > 100kb between adjacent hits. Initial peak bounds were then expanded to include any marginally significant hits (FDR *P* > 0.005 & ≤ 0.05) or fixed 21mer positions within 10kb of the peak interval bounds. All data processing, plotting, and analysis were accomplished in R 4.4.1 with the following packages: Biostrings v2.72.1 (Pagès et al. 2020), GenomicRanges v1.56.2 (Lawrence et al. 2013), ggplot2 v4.0.0 (Wickham 2016), and data.table v1.16.4 (Dowle and Srinivasan 2021).

### Total and allele-specific expression

Four tissue types: dormant buds, swelling buds, immature catkins, and immature pistillate flowers, were collected from mature grafted pecan trees (16–35 years old) of the cultivars ‘Apache’, ‘Mahan’, ‘Elliott’, ‘Lakota’, ‘Major’, and ‘Pawnee’ between January and April 2017. Immediately after sampling, tissues were flash frozen in liquid nitrogen to preserve RNA integrity. All trees were maintained at the USDA repository orchard and the National Pecan Advanced Clonal Test System in Somerville, Texas, USA. Total RNA was extracted using the Plant/Fungi Total RNA Purification Kit (Norgen Biotek, Ontario, Canada) following the manufacturer’s protocol. Stranded RNA-seq libraries were prepared with the NEBNext® Ultra™ RNA Library Prep Kit for Illumina® (NEB, USA), and unique index codes were incorporated to distinguish individual samples. Libraries were sequenced on an Illumina platform. Allele specific expression was assigned to either the Lakota-Mahan (dominant) or Lakota-Major (recessive) haplotype via competitive mapping (Lovell et al. 2018). In short, RNA-seq reads were aligned to the concatenated genome of both Lakota and Elliott haplotypes. Reads were identified as a) uniquely mapping to one haplotype (“[Haplotype]-ASE”) or equally mapping to both of the 1:1 orthologs (“[Haplotype]-shared”). Total expression counts for each genotype were the summed [Haplotype]-ASE values with the count of shared reads from the first haplotype.

Differential expression was assayed following standard protocols in DESeq2 (Love et al. 2014). Visualization of total transcript abundance used the variance stabilized transformation blind to the experimental design and significance was assayed with FDR-corrected likelihood ratio test *P*-values. Statistical inference of cis, strain and cis-by-tissue effects was conducted following (Westbrook et al. 2026). In short, we calculated library size factors from total ASE counts (see above) across both alleles, then applied these to ASE columns so that global bias in library size did not affect ASE bias (which should be the cis effect). With library sizes controlled for, we accomplished likelihood ratio tests in DESeq2 where: (1) “strain” effect was the difference between a full model and a reduced model without the main effect of strain and strain*tissue interactions were removed, (2) “cis-by-tissue” effect was the difference between the full model and the reduced model without the allele-tissue interaction terms, and (3) “cis” effect was the difference between the reduced model in #2 and one without the main effect of allele. In all cases, significance was calculated as FDR-corrected P-values of only the G-locus. We did not correct for genome-wide differential expression because we were a priori interested in only the G-locus. Significant effects had FDR-corrected *P*-values ≤ 0.05 and “strong” significant effects were *P* ≤ 0.005.

### Pangenome graph construction

The eight haplotype pangenome graph was constructed per chromosome with Minigraph-Cactus v3.0.0 (Hickey et al. 2023) using Lakota-Mahan as the primary reference. Graph processing was performed with the variation graph (vg) toolkit v1.68.0 (Garrison et al. 2018). Clipped chromosome graphs were combined with vg combine. Conversion from GFA to GBZ formats was handled with vg gbwt (Sirén and Paten 2022) and vg index. Presence-absence variation was identified by odgi PAV in 1kb windows based on the Lakota-Mahan assembly (odgi version v0.9.3-0-g497d046c, (Guarracino et al. 2022)). Regions of interest were extracted from the vcfwave VCF (Garrison et al. 2022) and processed with bcftools index, norm, sort, and +fixploidy version 1.22 (Danecek et al. 2021). Variants from the region of interest VCF were merged using truvari collapse (English et al. 2022). The merged and simplified VCF was used to construct a graph with vg construct and vg gbwt exclusively for visualization with SequenceTubeMap (Beyer et al. 2019).

### Short read library alignments to graph pangenome

K-mers were sampled from each short read library with kmc (options: -k29 -m200 -okff) (Kokot et al. 2017). Haplotypes in the graph were sampled with vg haplotypes based on k-mers present in each of the libraries (options: --include-reference --kmer-length 29 --subchain-length 10000 --kmer-input $KMC_OUT) (Sirén et al. 2024). Short read libraries were aligned to the clipped, haplotype-sampled subgraph with vg giraffe to produce a BAM file followed by sorting with samtools and deduplication by picard MarkDuplicates.

### Variant calling against graph pangenome

First, the Lakota-Mahan reference genome was split into 1Mb chunks. Base pair coverage was calculated with samtools mpileup 1.19.2 from a file list of BAM files (options: --min-BQ 20 -d 500). Variant calling was performed by varscan mpileup2cns v2.4.3 (options: --min-coverage 1 --min-reads2 0 --min-var-freq 0.001 --output-vcf 1). Using bcftools for all downstream processing, chunked VCF files were annotated and genotypes with read depth <8 were marked as missing. Last, 1Mb chunked VCFs were concatenated and provided as input for GWAS.

## Description of Supplementary Materials

**Supplementary Note 1** | **Genome assembly methods**. Complete assembly methods and basic statistical summaries for the seven new genome assemblies presented here.

**Supplementary Figure 1** | **Analysis of off-target GWAS peaks**.

Comparison of genotyping errors and their effects on GWAS peaks.

**Supplementary Figure 2** | **Allelic imbalance of all candidates and cultivars**. Variance-stabilized transformed total counts for all genes with reliable allele-specific expression (excluding UNK1, UNK2, CiCLE3) across the four tissues, two heterozygous cultivars and two G-locus haplotypes.

**Supplementary Figure 3** | **Runs of homozygosity in the Oaxaca reference genome**. The Illumina 2×150 polishing library was aligned against the Oaxaca HAP1 reference assembly. Runs of homozygous genotype calls (GATK) are visualized as red blocks of H0 = 0 along with a sliding window of heterozygosity genome wise.

**Supplementary Data 1** | **Short read resequencing panel metadata**. Phenotype, sample metadata, and library information for the 121 genotypes used in GWAS and the LD heatmap.

**Supplementary Data 2** | **Significant dichogamy-marker associations**. Position, GEMMA/LDAK statistics, and peak IDs for all marker-dichogamy associations with FDR-corrected P-values <= 0.05. These data correspond to the yellow, orange and red points for all five scans in Figure 1.

**Supplementary Data 3** | **Sliding windows for GWAS scans**. The most significant marker-trait association in all non-overlapping 100kb bins for five GWAS scans.

**Supplementary Data 4** | **Positions of dichogamy-associated kmers**. Data for all g-(found in all six *gg* libraries and never found in all six *GG* libraries) and *G*-21mers (opposite of g).

**Supplementary Data 5** | **Syntenic blocks between Lakota-Mahan and Lakota-Major G-locus sequences**. Alignment block file (.paf) generated by nucmer for the comparison of the two Lakota haplotypes plus a small buffer.

**Supplementary Data 6** | **G-locus gene positions**. Gene feature file (gff3)-formatted annotation for the G-locus in all eight genomes (field 3, source is labeled by genome; Oaxaca is synonymous with 87MX3).

**Supplementary Data 7** | **Tissue time course sample metadata**. Tissue, genotype and cultivar metadata for the transcript abundance experiment. The library field pairs with the counts matrix to allow for simple total RNA-seq transcript abundance analysis.

**Supplementary Data 8** | **Total RNA-seq counts**. Gene-by-library matrix containing uniquely mapping reads counted against the Lakota-Mahan reference genome and liftover annotation.

**Supplementary Data 9** | **Allele-specific and shared counts**.

Competitive mapping counts for orthologs of Lakota-Mahan and Lakota-Major. Like total counts, but with one row for each gene in the two annotations and two columns for each library: shared and unique.

## Acknowledgements

The work was also funded in part by U.S. Department of Agriculture National Institute of Food and Agriculture (SCRI-2016-51181-25408) “Coordinated Development of Genetic Tools for Pecan” and U.S. Department of Agriculture National Institute of Food and Agriculture (SCRI-2022-51181-38332) “Trees for the Future: Coordinated Development of Genetic Resources and Tools to Accelerate Breeding of Geographic Adapted Pecan Trees”, and by the U.S. Department of Agriculture – Agriculture Research Service National Programs through CRIS project 3091-21000-046-000-D (Crop Germplasm Research Unit, TX). The agency was not involved in the study design, collection, analysis, interpretation of data and the writing of this article. This article reports on the results of the research only. A.H. is supported by an NSF IOS-PGRP CAREER award 2239530. JTL, AMH, and CMM were supported by the Center for Bioenergy Innovation, which is supported by the U.S. Department of Energy, Office of Science, Biological and Environmental Research under Contract Number ERKP886. This manuscript was improved by thoughtful advice from J. Groh, and K Greenham.

## Data availability

Reference genome assemblies will all be made available on the Pecan Toolbox genomic toolset (https://pecantoolbox.nmsu.edu/genomes.html) upon publication. In the meantime, contact the authors for pre-publication access to the assemblies. All other data required to reproduce analyses presented can be found in the supplementary material.

