## Supplementary Information for "Dissecting the regulatory and genomic drivers of the dichogamy determining G-locus in pecan"

**Running title:** Comparative genomics of dichogamy in pecan

**John T. Lovell[1,2*], Hormat Rhein[3], Avinash Sreedasyam[1,2], Nolan Bentley[4], Patricia Klein[5], Warren Chatwin[6], Lillian Padgitt-Cobb[1], Avril M. Harder[1], Paul Grabowski[1], Sarah B. Carey[1], Alex Harkess[1], Jerry Jenkins[1], Chloee M. McLaughlin[1], Christopher Plott[1], Joanna Rifkin[1,2], Joe Song[3], Jenell Webber[1], Melissa Williams[1], Jane Grimwood[1], Jeremy Schmutz[1,2], L. J. Grauke[6], Xinwang Wang[5*] & Jennifer J. Randall[3*]**

[1] Genome Sequencing Center, HudsonAlpha Institute for Biotechnology, Huntsville, AL 35806, USA
[2] U.S. Department of Energy Joint Genome Institute, Lawrence Berkeley National Laboratory, Berkeley, CA 94720, USA
[3] New Mexico State University, Las Cruces, NM 88003, USA
[4] University of Texas at Austin, Austin, TX, USA
[5] USDA-ARS, Crop Germplasm Research Unit, College Station, TX, USA

#### Supplementary Figures

##### Supplementary Figure 1 | Pseudoheterozygous calls underlie GWAS peaks when aligning to a linear reference containing a recessive G-locus haplotype.

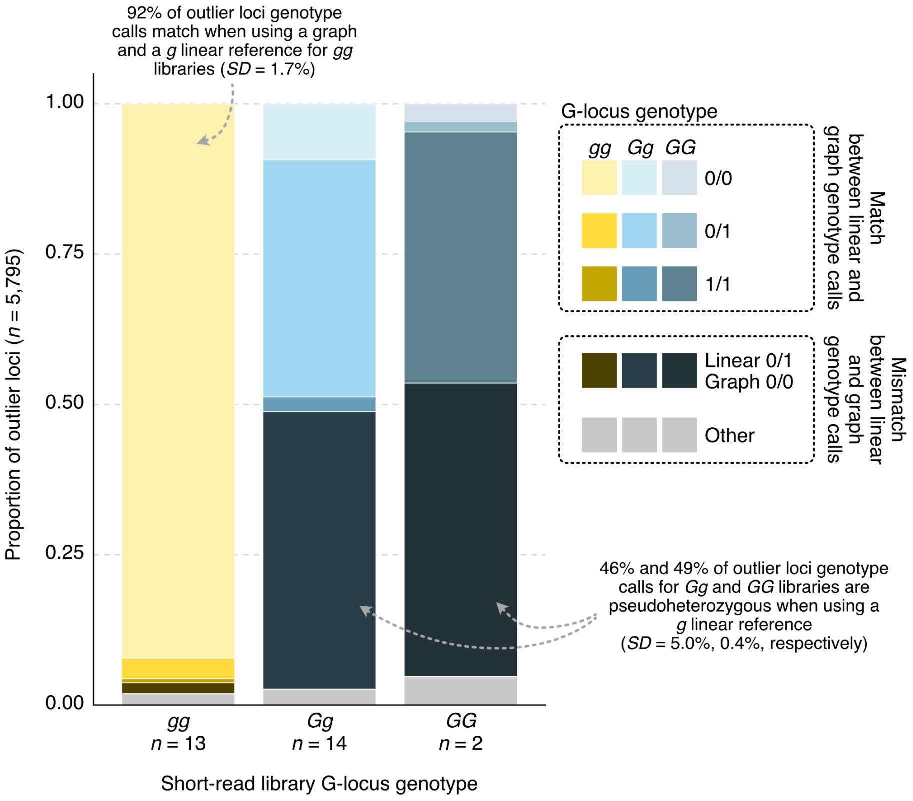

**Supplementary Figure 1 | Pseudoheterozygous calls underlie GWAS peaks when aligning to a linear reference containing a recessive G-locus haplotype.** Variants were called for 30 libraries aligned to both the Lakota-Major (*g*) reference and a graph reference resource built with Minigraph-Cactus, setting the Lakota-Major assembly as the primary reference. For the 5,975 SNPs identified as significant in the linear-based GWAS result, genotypes were compared at each site between the linear- and graph-based calls for each library. On average, 97% (*SD* = 1.3%) of these calls were identical between the two methods for *gg* libraries, with the vast majority of these sites called as reference homozygous genotypes (mean = 92% of sites, *SD* = 1.7%). However, for *Gg* and *GG* libraries, only 51% and 47% of variant calls were concordant between the two methods, respectively (*SD* = 4.1% and 0.06%). Most of the discordance in genotype calls comprises apparent pseudoheterozygous calls that result from poor alignment of *G* haplotype-origin short reads to a *g* linear reference. Lighter colors in the stacked bars indicate mean call agreement frequencies between linear- and graph-based methods, whereas darker colors and grey indicate mean mismatch frequencies.

### Supplementary Figure 2 | Allelic imbalance in F1s

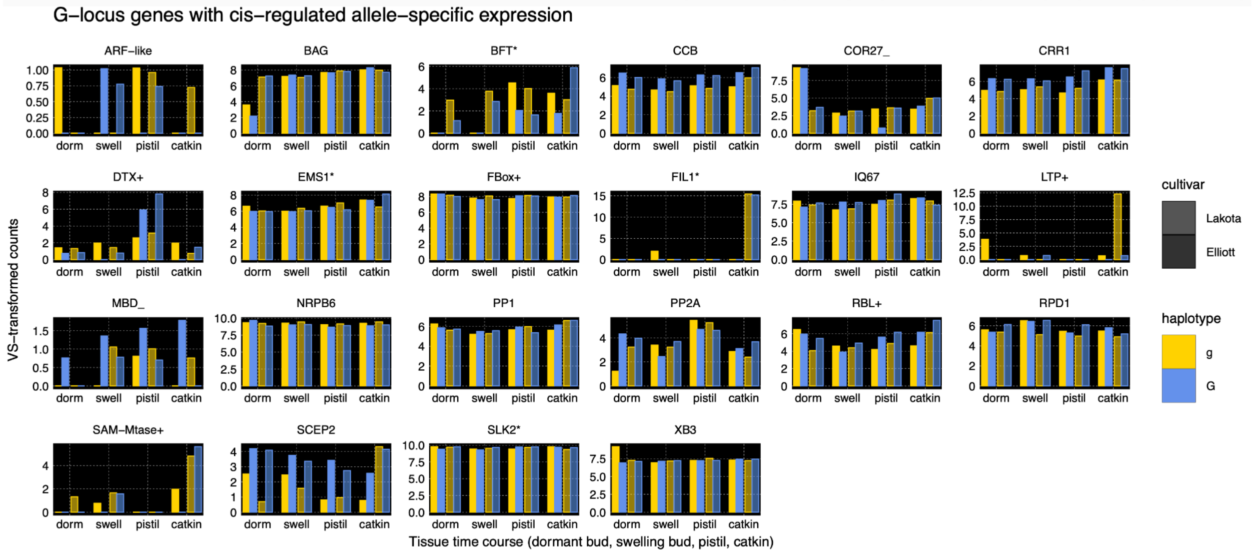

**Supplementary Figure 2 | Allelic imbalance in Lakota and Elliott**. Variance-stabilized transformed total counts for all genes with reliable allele-specific expression (excluding *UNK1*, *UNK2*, *CiCLE3*) across the four tissues, two heterozygous cultivars and two G-locus haplotypes.

### Supplementary Figure 3 | Runs of homozygosity in the Oaxaca genome

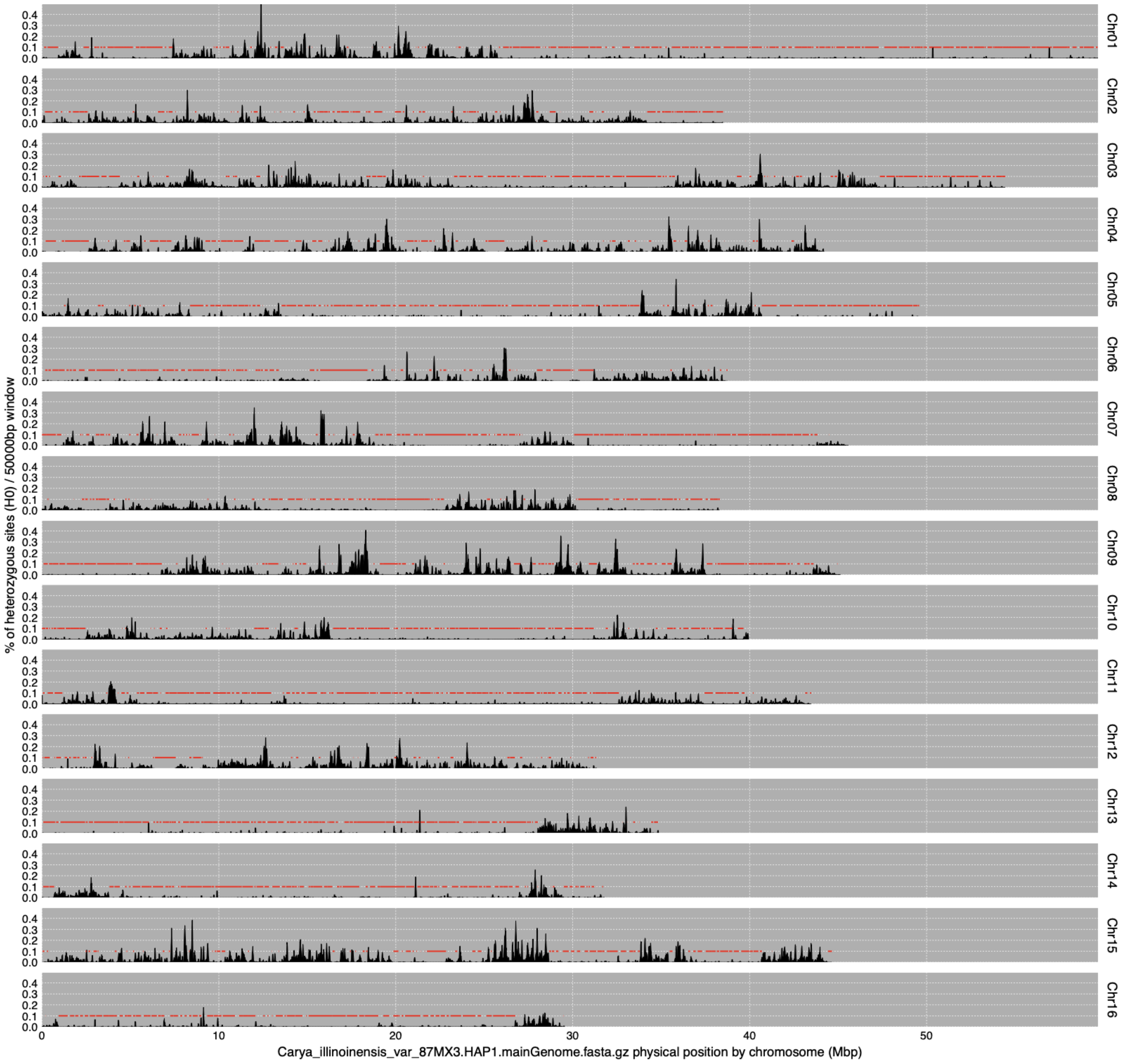

**Supplementary Figure 3 | Runs of homozygosity in the Oaxaca genome.** Variants called by GATK from the PCR-free polishing Illumina 2x150 library of the Oaxaca (87Mx3-2.11) genotype was aligned to the Oaxaca HAP1 reference. Heterozygous SNPs were parsed, and their density was plotted in 40kb-overlapping 50kb intervals (black areas). Runs of regions with reference homozygote calls (ROHs) are flagged as red segments.

### Supplementary Notes

#### Supplementary Note 1 | Detailed genome assembly methods

**Carya illinoinensis var. 87MX3 (HAP2) version 3.0**

- Main genome scaffold total: 17
- Main genome contig total: 86
- Main genome scaffold sequence total: 659.0 MB
- Main genome contig sequence total: 658.3 MB (-> 0.1% gap)
- Main genome scaffold L/N50: 7/42.9 MB
- Main genome contig L/N50: 17/15.9 MB
- Number of scaffolds > 50 KB: 17
- % main genome in scaffolds > 50 KB: 100.0%

Main assembly consisted of 38.56x of single haplotype CCS PACBIO coverage (6,132 bp average read size), and was assembled using HiFiAsm+HIC [CCS Library: PFBW; HIC Library: IPHL] and the resulting sequence was polished using RACON. Misjoins in the assembly were identified using HIC data. There were no misjoins identified in the polished assembly. Contigs for both haplotypes were then oriented, ordered, and joined into chromosomes using the JUICER pipeline (HiC Library: IPHL). Contigs terminating in significant telomeric sequence were properly oriented in the assembly. A total of 90 joins were applied to the assembly to form the final assembly consisting of 16 chromosomes. A total of 99.98% of the assembled sequence is contained in the chromosomes. Adjacent alternative haplotypes were identified on the joined contig set. Althap regions were collapsed using the longest common substring between the two haplotypes. 21 adjacent alternative haplotypes were collapsed in the assembly. All homopolymer regions longer than 500bp were evaluated as potential sequencing artifacts using aligned CCS depth, and no regions were removed. The chloroplast genome was assembled using OatK (https://github.com/c-zhou/oatk), however there was insufficient evidence to assemble the mitochondrial genome. Chromosomes were numbered using the V1 C. illinoinensis var. Pawnee release. The resulting sequence was screened for retained vector and/or contaminants.

Misphased regions were identified by aligning the HiC data to the combined HAP1/HAP2 chromosomes and looking for regions of the chromosomes that need to be swapped. The regions that were swapped are given below:

- Chr06 HAP1 32,206,149 32,409,321 HAP2 30,171,755 30,397,619
- Chr06 HAP1 31,190,542 31,400,539 HAP2 29,240,294 29,425,294
- Chr06 HAP1 34,811,850 35,615,334 HAP2 32,693,912 33,488,558
- Chr09 HAP1 10,433,920 10,888,322 HAP2 10,476,557 10,910,410

Homozygous SNPs and INDELs were corrected in the release sequence using ~55.5x (library code IIMT/IIMU) of Illumina reads (2x150, 400bp insert). The table below provides the summary statistics of this correction.

| Case | Het_SNPs | Hom_SNPs | Het_INDELs | Hom_INDELs | Callable Bases |
|---|---|---|---|---|---|
| Pre SNP/INDEL fixing | 2,125,137 | 346 | 209,165 | 4,944 | 641,200,656 |
| Post SNP/INDEL fixing | 2,125,169 | 45 | 209,422 | 24 | 641,201,369 |

**Carya illinoinensis var. Elliot (HAP1) version 2.0**

- Main genome scaffold total: 17
- Main genome contig total: 26
- Main genome scaffold sequence total: 680.9 MB
- Main genome contig sequence total: 680.8 MB (-> 0.0% gap)
- Main genome scaffold L/N50: 7/45.1 MB
- Main genome contig L/N50: 8/38.0 MB
- Number of scaffolds > 50 KB: 17
- % main genome in scaffolds > 50 KB: 100.0%

Main assembly consisted of 114.86x of single haplotype CCS PACBIO coverage (14,763 bp average read size), and was assembled using HiFiAsm+HIC [CCS Library: PFBX; HIC Library: ITIR] and the resulting sequence was polished using RACON. Misjoins in the assembly were identified using HIC data. There were no misjoins identified in the polished assembly. Contigs for both haplotypes were then oriented, ordered, and joined into chromosomes using the JUICER pipeline (HiC Library: ITIR). Contigs terminating in significant telomeric sequence were properly oriented in the assembly. A total of 12 joins were applied to the assembly to form the final assembly consisting of 16 chromosomes. A total of 99.991% of the assembled sequence is contained in the chromosomes. Adjacent alternative haplotypes were identified on the joined contig set. Althap regions were collapsed using the longest common substring between the two haplotypes. 3 adjacent alternative haplotypes were collapsed in the assembly. All homopolymer regions longer than 500bp were evaluated as potential sequencing artifacts using aligned CCS depth, and no regions were removed. The chloroplast and mitochondrial genome was assembled using OatK (https://github.com/c-zhou/oatk). Chromosomes were numbered using the V1 C. illinoinensis var. Pawnee release. The resulting sequence was screened for retained vector and/or contaminants.

Misphased regions were identified by aligning the HiC data to the combined HAP1/HAP2 chromosomes and looking for regions of the chromosomes that need to be swapped. The regions that were swapped are given below:

| Chromosome | Haplotype | start | end | Haplotype | start | end |
|---|---|---|---|---|---|---|
| • Chr01 | HAP1 | 38,883,313 | 40,040,900 | HAP2 | 38,535,526 | 39,674,575 |

Homozygous SNPs and INDELs were corrected in the release sequence using ~55.0x (library code IPZM) of Illumina reads (2x150, 400bp insert). The table below provides the summary statistics of this correction.

| Case | Het_SNPs | Hom_SNPs | Het_INDELs | Hom_INDELs | Callable Bases |
|---|---|---|---|---|---|
| Pre SNP/INDEL fixing | 4,614,512 | 348 | 478,606 | 7,239 | 667,974,703 |
| Post SNP/INDEL fixing | 4,614,691 | 33 | 478,930 | 36 | 667,972,201 |

**Carya illinoinensis var. Elliot (HAP2) version 2.0**

- Main genome scaffold total: 16
- Main genome contig total: 27
- Main genome scaffold sequence total: 665.1 MB
- Main genome contig sequence total: 665.0 MB (-> 0.0% gap)
- Main genome scaffold L/N50: 7/43.9 MB
- Main genome contig L/N50: 9/33.7 MB
- Number of scaffolds > 50 KB: 16
- % main genome in scaffolds > 50 KB: 100.0%

Main assembly consisted of 114.86x of single haplotype CCS PACBIO coverage (14,763 bp average read size), and was assembled using HiFiAsm+HIC [CCS Library: PFBX; HIC Library: ITIR] and the resulting sequence was polished using RACON. Misjoins in the assembly were identified using HIC data. There were no misjoins identified in the polished assembly.Contigs for both haplotypes were then oriented, ordered, and joined into chromosomes using the JUICER pipeline (HiC Library: ITIR). Contigs terminating in significant telomeric sequence were properly oriented in the assembly. A total of 14 joins were applied to the assembly to form the final assembly consisting of 16 chromosomes. A total of 100% of the assembled sequence is contained in the chromosomes. Adjacent alternative haplotypes were identified on the joined contig set. Althap regions were collapsed using the longest common substring between the two haplotypes. 3 adjacent alternative haplotypes were collapsed in the assembly. All homopolymer regions longer than 500bp were evaluated as potential sequencing artifacts using aligned CCS depth, and no regions were removed. The chloroplast and mitochondrial genome was assembled using OatK (https://github.com/c-zhou/oatk). Chromosomes were numbered using the V1 C. illinoinensis var. Pawnee release. The resulting sequence was screened for retained vector and/or contaminants.

Misphased regions were identified by aligning the HiC data to the combined HAP1/HAP2 chromosomes and looking for regions of the chromosomes that need to be swapped. The regions that were swapped are given below:

| Chromosome | Haplotype | start | end | Haplotype | start | end |
|---|---|---|---|---|---|---|
| Chr01 | HAP1 | 38,883,313 | 40,040,900 | HAP2 | 38,535,526 | 39,674,575 |

Homozygous SNPs and INDELs were corrected in the release sequence using ~56.9x (library code IPZM) of Illumina reads (2x150, 400bp insert). The table below provides the summary statistics of this correction.

| Case | Het_SNPs | Hom_SNPs | Het_INDELs | Hom_INDELs | Callable Bases |
|---|---|---|---|---|---|
| Pre SNP/INDEL fixing | 4,595,589 | 316 | 473,350 | 6,574 | 652,030,430 |
| Post SNP/INDEL fixing | 4,595,670 | 36 | 473,666 | 51 | 652,030,804 |

**Carya illinoinensis var. Pawnee (Mohawk) version 2.0**

- Main genome scaffold total: 16
- Main genome contig total: 78
- Main genome scaffold sequence total: 671.7 MB
- Main genome contig sequence total: 671.1 MB (-> 0.1% gap)
- Main genome scaffold L/N50: 7/44.4 MB
- Main genome contig L/N50: 16/15.2 MB
- Number of scaffolds > 50 KB: 16
- % main genome in scaffolds > 50 KB: 100.0%

The V3 assembly was phased into parental chromosomes (Mohawk and SHG) by aligning the Illumina reads from the libraries IWWM (Mohawk) & IYBA (SHG), and examining the homozygous SNP counts on the chromosome pairs. No modifications were made to any of the sequences, aside from the re-partitioning.

Main assembly consisted of 52.12x of single haplotype CCS PACBIO coverage (20,869 bp average read size), and was assembled using HiFiAsm+HIC [CCS Library: PBXT,PBXU; HIC Library: IUSF] and the resulting sequence was polished using RACON. Misjoins in the assembly were identified using HIC data. There were no misjoins identified in the polished assembly. Contigs for both haplotypes were then oriented, ordered, and joined into chromosomes using the JUICER pipeline (HiC Library: IUSF). Contigs terminating in significant telomeric sequence were properly oriented in the assembly. A total of 64 joins were applied to the assembly to form the final assembly consisting of 16 chromosomes. A total of 100.0% of the assembled sequence is contained in the chromosomes. Adjacent alternative haplotypes were identified on the joined contig set. Althap regions were collapsed using the longest common substring between the two haplotypes. 5 adjacent alternative haplotypes were collapsed in the assembly. All homopolymer regions longer than 500bp were evaluated as potential sequencing artifacts using aligned CCS depth, and 1 region was removed. The chloroplast and mitochondrial genome was assembled using OatK (https://github.com/c-zhou/oatk). Chromosomes were numbered using the V1 C. illinoinensis var. Pawnee release. The resulting sequence was screened for retained vector and/or contaminants.

Misphased regions were identified by aligning the HiC data to the combined HAP1/HAP2 chromosomes and looking for regions of the chromosomes that need to be swapped. The regions that were swapped are given below:

| Chromosome | Haplotype | start | end | Haplotype | start | end |
|---|---|---|---|---|---|---|
| Chr04 | HAP1 | 14,184,985 | 29,168,910 | HAP2 | 14,078,600 | 25,636,577 |

Homozygous SNPs and INDELs were corrected in the release sequence using ~52.3x (library code JAED) of Illumina reads (2x150, 400bp insert). The table below provides the summary statistics of this correction.

| Case | Het_SNPs | Hom_SNPs | Het_INDELs | Hom_INDELs | Callable Bases |
|---|---|---|---|---|---|
| Pre SNP/INDEL fixing | 4,910,630 | 341 | 505,179 | 6,826 | 658,904,498 |
| Post SNP/INDEL fixing | 4,911,246 | 68 | 507,780 | 122 | 658,898,921 |

**Carya illinoinensis var. Pawnee (SHG) version 2.0**

- Main genome scaffold total: 19
- Main genome contig total: 78
- Main genome scaffold sequence total: 665.6 MB
- Main genome contig sequence total: 665.0 MB (-> 0.1% gap)
- Main genome scaffold L/N50: 7/44.3 MB
- Main genome contig L/N50: 15/13.9 MB
- Number of scaffolds > 50 KB: 19
- % main genome in scaffolds > 50 KB: 100.0%

The V3 assembly was phased into parental chromosomes (Mohawk and SHG) by aligning the Illumina reads from the libraries IWWM (Mohawk) & IYBA (SHG), and examining the homozygous SNP counts on the chromosome pairs. No modifications were made to any of the sequences, aside from the re-partitioning.

Main assembly consisted of 52.12x of single haplotype CCS PACBIO coverage (20,869 bp average read size), and was assembled using HiFiAsm+HIC [CCS Library: PBXT,PBXU; HIC Library: IUSF] and the resulting sequence was polished using RACON. Misjoins in the assembly were identified using HIC data. There were no misjoins identified in the polished assembly. Contigs for both haplotypes were then oriented, ordered, and joined into chromosomes using the JUICER pipeline (HiC Library: IUSF). Contigs terminating in significant telomeric sequence were properly oriented in the assembly. A total of 64 joins were applied to the assembly to form the final assembly consisting of 16 chromosomes. A total of 99.91% of the assembled sequence is contained in the chromosomes. Adjacent alternative haplotypes were identified on the joined contig set. Althap regions were collapsed using the longest common substring between the two haplotypes. 3 adjacent alternative haplotypes were collapsed in the assembly. All homopolymer regions longer than 500bp were evaluated as potential sequencing artifacts using aligned CCS depth, and no regions were removed. The chloroplast and mitochondrial genome was assembled using OatK (https://github.com/c-zhou/oatk). Chromosomes were numbered using the V1 C. illinoinensis var. Pawnee release. The resulting sequence was screened for retained vector and/or contaminants.

Misphased regions were identified by aligning the HiC data to the combined HAP1/HAP2 chromosomes and looking for regions of the chromosomes that need to be swapped. The regions that were swapped are given below:

| Chromosome | Haplotype | start | end | Haplotype | start | end |
|---|---|---|---|---|---|---|
| • Chr04 | HAP1 | 14,184,985 | 29,168,910 | HAP2 | 14,078,600 | 25,636,577 |

Homozygous SNPs and INDELs were corrected in the release sequence using ~52.4x (library code JAED) of Illumina reads (2x150, 400bp insert). The table below provides the summary statistics of this correction.

| Case | Het_SNPs | Hom_SNPs | Het_INDELs | Hom_INDELs | Callable Bases |
|---|---|---|---|---|---|
| Pre SNP/INDEL fixing | 4,943,444 | 419 | 508,333 | 11,834 | 650,686,347 |
| Post SNP/INDEL fixing | 4,944,226 | 83 | 510,922 | 100 | 650,681,271 |

**Carya illinoinensis var. Lakota (Mahan) version 4.0**

- Main genome scaffold total: 17
- Main genome contig total: 27
- Main genome scaffold sequence total: 676.3 MB
- Main genome contig sequence total: 676.2 MB (-> 0.0% gap)
- Main genome scaffold L/N50: 7/45.2 MB
- Main genome contig L/N50: 9/34.3 MB
- Number of scaffolds > 50 KB: 17
- % main genome in scaffolds > 50 KB: 100.0%

After releasing the V3, it was discovered that Chr12 was not phased properly. Chr12 was switched between Lakota_Major and Lakota_Mahan and verified by calling snps using the IYBQ Major resequencing library. The V3 assembly was phased into parental chromosomes (Mohawk and SHG) by aligning the Illumina reads from the libraries IXWF (Mahan) & IYBQ (Major), and examining the homozygous SNP counts on the chromosome pairs. The partitioning of HAP1/HAP2 into Mahan and Major from the V2 release is given in the table below. No modifications were made to any of the sequences, aside from the re-partitioning.

Main assembly consisted of 67.25x of single haplotype CCS PACBIO coverage (13,903 bp average read size), and was assembled using HiFiAsm+HIC [CCS Library: PFBY; HIC Library: IPHK] and the resulting sequence was polished using RACON. Misjoins in the assembly were identified using HIC data. There were no misjoins identified in the polished assembly. Contigs for both haplotypes were then oriented, ordered, and joined into chromosomes using the JUICER pipeline (HiC Library: IPHK). Contigs terminating in significant telomeric sequence were properly oriented in the assembly. A total of 18 joins were applied to the assembly to form the final assembly consisting of 16 chromosomes. A total of 99.992% of the assembled sequence is contained in the chromosomes. Adjacent alternative haplotypes were identified on the joined contig set. Althap regions were collapsed using the longest common substring between the two haplotypes. 5 adjacent alternative haplotypes were collapsed in the assembly. All homopolymer regions longer than 500bp were evaluated as potential sequencing artifacts using aligned CCS depth, and no regions were removed. The chloroplast and mitochondrial genome was assembled using OatK (https://github.com/c-zhou/oatk). Chromosomes were numbered using the V1 C. illinoinensis var. Pawnee release. The resulting sequence was screened for retained vector and/or contaminants.

Misphased regions were identified by aligning the HiC data to the combined HAP1/HAP2 chromosomes and looking for regions of the chromosomes that need to be swapped. The regions that were swapped are given below:

| Chromosome | Haplotype | start | end | Haplotype | start | end |
|---|---|---|---|---|---|---|
| Chr04 | HAP1 | 36,978,115 | 45,208,478 | HAP2 | 36,441,915 | 44,786,744 |

Homozygous SNPs and INDELs were corrected in the release sequence using ~52.5x (library code IKFW) of Illumina reads (2x150, 400bp insert). The table below provides the summary statistics of this correction.

| Case | Het_SNPs | Hom_SNPs | Het_INDELs | Hom_INDELs | Callable Bases |
|---|---|---|---|---|---|
| Pre SNP/INDEL fixing | 5,052,761 | 302 | 525,871 | 6,658 | 659,008,660 |
| Post SNP/INDEL fixing | 5,052,942 | 48 | 526,230 | 73 | 659,001,927 |

**Carya illinoinensis var. Lakota (Major) version 4.0**

- Main genome scaffold total: 16
- Main genome contig total: 28
- Main genome scaffold sequence total: 666.7 MB
- Main genome contig sequence total: 666.5 MB (-> 0.0% gap)
- Main genome scaffold L/N50: 7/44.3 MB
- Main genome contig L/N50: 8/37.3 MB
- Number of scaffolds > 50 KB: 16
- % main genome in scaffolds > 50 KB: 100.0%

After releasing the V3, it was discovered that Chr12 was not phased properly. Chr12 was switched between Lakota_Major and Lakota_Mahan and verified by calling snps using the IYBQ Major resequencing library. The V3 assembly was phased into parental chromosomes (Mohawk and SHG) by aligning the Illumina reads from the libraries IXWF (Mahan) & IYBQ (Major), and examining the homozygous SNP counts on the chromosome pairs. The partitioning of HAP1/HAP2 into Mahan and Major from the V2 release is given in the table below. No modifications were made to any of the sequences, aside from the re-partitioning. Main assembly consisted of 67.25x of single haplotype CCS PACBIO coverage (13,903 bp average read size), and was assembled using HiFiAsm+HIC

[CCS Library: PFBY; HIC Library: IPHK] and the resulting sequence was polished using RACON. Misjoins in the assembly were identified using HIC data. There were no misjoins identified in the polished assembly.

Contigs for both haplotypes were then oriented, ordered, and joined into chromosomes using the JUICER pipeline (HiC Library: IPHK). Contigs terminating in significant telomeric sequence were properly oriented in the assembly. A total of 12 joins were applied to the assembly to form the final assembly consisting of 16 chromosomes. A total of 100% of the assembled sequence is contained in the chromosomes. Adjacent alternative haplotypes were identified on the joined contig set. Althap regions were collapsed using the longest common substring between the two haplotypes. 3 adjacent alternative haplotypes were collapsed in the assembly. All homopolymer regions longer than 500bp were evaluated as potential sequencing artifacts using aligned CCS depth, and no regions were removed. The chloroplast and mitochondrial genome was assembled using OatK (https://github.com/c-zhou/oatk). Chromosomes were numbered using the V1 C. illinoinensis var. Pawnee release. The resulting sequence was screened for retained vector and/or contaminants.

Misphased regions were identified by aligning the HiC data to the combined HAP1/HAP2 chromosomes and looking for regions of the chromosomes that need to be swapped. The regions that were swapped are given below:

| Chromosome | Haplotype | start | end | Haplotype | start | end |
|---|---|---|---|---|---|---|
| Chr04 | HAP1 | 36,978,115 | 45,208,478 | HAP2 | 36,441,915 | 44,786,744 |

Homozygous SNPs and INDELs were corrected in the release sequence using ~55.4x (library code IKFW) of Illumina reads (2x150, 400bp insert). The table below provides the summary statistics of this correction.

| Case | Het_SNPs | Hom_SNPs | Het_INDELs | Hom_INDELs | Callable Bases |
|---|---|---|---|---|---|
| Pre SNP/INDEL fixing | 5,043,631 | 300 | 530,257 | 6,715 | 643,675,071 |
| Post SNP/INDEL fixing | 5,043,768 | 52 | 530,665 | 52 | 643,675,636 |
